# Relevance of Auditory Errors Decreases When Errors Are Introduced Suddenly

**DOI:** 10.1101/2021.08.09.455646

**Authors:** Sara-Ching Chao, Ayoub Daliri

## Abstract

**Purpose:** When the speech motor system encounters errors, it generates adaptive responses to compensate for the errors. We previously showed that adaptive responses to task-irrelevant errors are significantly smaller than responses to task-relevant errors when errors are introduced gradually. The current study aimed to examine responses to task-irrelevant and task-relevant errors when errors are introduced suddenly.

**Method:** We used an adaptation paradigm in which participants experienced task-relevant errors (induced by formant-shift perturbations) and task-irrelevant errors (induced by formant-clamp perturbations). For one group of participants (*N* = 30), we applied the perturbations gradually. The second group of participants (*N* = 30) received the perturbations suddenly. We designed the perturbations based on participant-specific vowel configurations such that a participant’s first and second formants of /ε/ were perturbed toward their /æ/. To estimate adaptive responses, we measured formant changes (within 0–100 ms of the vowel onset) in response to the formant perturbations.

**Results:** We found that (1) the difference between adaptive responses to formant-shift and formant-clamp perturbations was the smallest for the suddenly introduced perturbations, and (2) responses to formant-shift perturbations positively correlated with responses to formant-clamp perturbations for the suddenly (but not gradually) introduced perturbations.

**Conclusions:** These results showed that the speech motor system responds to task-relevant errors and task-irrelevant errors more differently when errors are introduced gradually than suddenly. Overall, the speech motor system evaluates the relevance of errors and uses its evaluation to modulate its adaptive responses to errors.

---

Relevance of Auditory Errors Decreases When Errors Are Introduced Suddenly Speech monitoring plays a fundamental role in speech acquisition during childhood and ensures speech production accuracy during adulthood (Callan et al., 2000; Guenther, 2016). As the speech motor system (SMS) prepares speech movements, it also predicts somatosensory and auditory outcomes of the movements (Daliri, 2021; Daliri et al., 2014; Guenther, 2016; Hickok, 2012; Houde & Nagarajan, 2011; Parrell et al., 2019). The SMS then executes the planned movements and monitors the sensory outcomes of the movements to ensure that the movements are executed accurately. For this purpose, the SMS compares the predicted sensory outcomes of the movements with the actual sensory feedback from the movements. During this speech monitoring process, the SMS may find a mismatch between the predicted outcome and sensory feedback (i.e., *prediction error*). The SMS uses the sensory prediction error to modify (*adapt*) its motor programs so that future speech movements more accurately reach the desired sensory targets (Daliri, 2021; Kearney et al., 2020). Thus, error detection and error adaptation are crucial processes that ensure the accuracy of speech movements.

Prediction errors typically occur due to structural changes in speech articulators (e.g., rapid developmental changes during childhood or structural changes during the aging process) or sensory feedback (e.g., hearing impairment) (Daliri et al., 2013; Perkell, 2012; Perkell et al., 1992; Pittman et al., 2018; Vorperian et al., 2005, 2009). One could generate prediction errors in the laboratory setting by experimentally manipulating somatosensory feedback (Daliri et al., 2013; Lametti et al., 2012; Tremblay et al., 2003) or auditory feedback (Abur et al., 2018; Ballard et al., 2018; Daliri et al., 2017; Daliri & Max, 2018; Houde & Jordan, 1998; Kim et al., 2020; Kothare et al., 2020; Scott et al., 2020; Stepp et al., 2017). Due to practical reasons and the importance of auditory feedback for speech production, many studies have used auditory perturbations to induce auditory prediction errors during speaking (for a review, see Fuchs et al., 2019). In auditory perturbation paradigms, acoustic characteristics of a speaker’s speech, such as formants, are shifted in near real-time and played back to the speaker (e.g., the speaker produces the word “bed” but hears a word that sounds like “bad”). The *formant-shift* procedure results in an experimentally generated auditory prediction error—a mismatch between the speaker’s expected auditory feedback (“bed”) and actual auditory feedback (“bad”). In response to repeated exposure to prediction errors, the speaker gradually adapts his/her motor commands to reduce the error. Examining adaptive responses provides essential insights into the error detection and error adaptation processes that the SMS uses to integrate prediction errors.

Previous studies have shown that characteristics of prediction errors contribute to the magnitude of adaptive responses (Daliri & Dittman, 2019; Kothare et al., 2020; MacDonald et al., 2010; Max & Maffett, 2015; Mitsuya et al., 2017). For example, adaptive responses to substantially large errors (MacDonald et al., 2010) or errors that are introduced with a delay (Max & Maffett, 2015; Mitsuya et al., 2017) are minimal or entirely absent. We have interpreted such findings as evidence that the SMS evaluates errors and responds less to less task-relevant errors (Daliri & Dittman, 2019). We recently developed a novel experimental procedure (i.e., *formant-clamp*) to directly manipulate the relevance of auditory errors. In the formant-clamp procedure, the induced error does not correspond with the speaker’s speech (i.e., task-irrelevant error), and the speaker cannot generate adaptive responses to reduce the experienced error. We found that adaptive responses to formant-clamp perturbations were significantly smaller than responses to formant-shift perturbations (Daliri & Dittman, 2019). In that study, we applied the perturbations gradually; thus, the SMS was exposed to small errors that incrementally increased over several trials, allowing the SMS to develop adaptive responses to the experienced errors. Studies of limb motor learning suggest that different neural processes may prepare adaptive responses to gradually and suddenly introduced errors (Doya, 2000; Kim et al., 2020; Smith & Shadmehr, 2005; Venkatakrishnan et al., 2011; Werner et al., 2010). By examining adaptive responses to formant-shift perturbations, previous studies have shown that the SMS generates similar adaptive responses to gradually and suddenly introduced auditory errors (Kearney et al., 2020; Kim et al., 2020; MacDonald et al., 2010). However, it remained unclear how the SMS would evaluate and respond to suddenly introduced formant-clamp perturbations.

The primary goal of the present study is to determine how the SMS evaluates and responds to task-relevant vs. task-irrelevant errors that are introduced suddenly or gradually. We used an auditory-motor adaptation paradigm similar to our previous study (Daliri & Dittman, 2019), in which we applied formant-shift and formant-clamp perturbations. We used gradually introduced perturbations in our previous study (Daliri & Dittman, 2019); thus, we used suddenly introduced formant-shift and formant-clamp perturbations in the current study. We designed the perturbations based on participant-specific vowel configurations such that a participant’s first and second formants of /ε/ were perturbed toward their /æ/. The magnitude of the perturbation was 80% of the participant-specific ε–æ distance. We measured formant changes within 0–100 ms from the vowel onset to estimate adaptive responses. We previously showed that these early responses are more appropriately suited to study adaptive changes (Daliri, 2021). We also re-analyzed the published data from participants in our previous study in which we applied gradual perturbations (Daliri & Dittman, 2019). Overall, we compared adaptive responses to suddenly introduced formant-shift and formant-clamp perturbations (data from the current study) with responses to gradually introduced formant-shift and formant-clamp perturbations (data from Daliri & Dittman, 2019). Suddenly introduced perturbations are more noticeable than gradually introduced perturbations; thus, at least initially, the SMS may evaluate suddenly introduced errors as less task-relevant errors. We hypothesized that the SMS responds to task-relevant errors (induced by formant-shift perturbations) and task-irrelevant errors (induced by formant-clamp perturbations) more differently when errors are introduced gradually than suddenly.

## Method

### Participants

In this study, we included data from two groups of participants with similar participant characteristics: (1) a new group of participants and (2) a group of participants from our published study (Daliri & Dittman, 2019). The first group included 30 healthy adult participants (age: *M* = 24.4 years, *SD* = 5.3 years; age range: 20.7–45.8 years). Most participants were recruited from a pool of undergraduate students at Arizona State University. All participants met the following inclusion criteria: (1) being a native speaker of American English, (2) not having a current or a history of speech-language, psychological, and neurological disorders (self-reported), (3) not taking medications that influence the nervous system, and (4) having normal binaural hearing thresholds (≤20 dB HL at all octave frequencies of 250–8000 Hz) as tested with a standard pure-tone hearing screening (ASHA, 1997). The study protocols were approved by the Institutional Review Board of Arizona State University, and participants signed a written consent form before the experiment.

The second group included 30 healthy adult participants (age: *M* = 23.7 years, *SD* = 3.7 years; age range: 18.5–32.9 years). We have already published the data from these participants (Daliri & Dittman, 2019). However, we re-analyzed their data and included the analysis in the present study to compare adaptive responses to suddenly introduced (current study) with gradually introduced (previous study) auditory perturbations. More information about these participants can be found in our previous study (Daliri & Dittman, 2019).

### Apparatus

As shown in Figure 1A, we used the same experimental apparatus as our previous study (Daliri & Dittman, 2019). Participants sat comfortably in a chair in front of a computer monitor. We recorded speech using a microphone (Shure SM58), placed 15 cm away from the participant’s mouth at a 45° angle. The microphone signal was amplified (ART Tubeopto 8) and fed into an audio interface (MOTU Ultralite Mk3 hybrid). This audio interface was connected to a computer that controlled all aspects of the experiment, such as presenting visual stimuli to the participant and manipulating auditory feedback. The audio interface’s output was amplified (Eurorack Pro Rx1602) and binaurally played back to the participant via insert earphones (ER-1, Etymotic Research Inc.). The amplification level of the microphone and headphones amplifiers was adjusted so that the headphones signals were played back at 5 dB higher than the microphone signal (Daliri & Max, 2015, 2016; McGuffin et al., 2020; Merrikhi et al., 2018).

**Figure 1.**
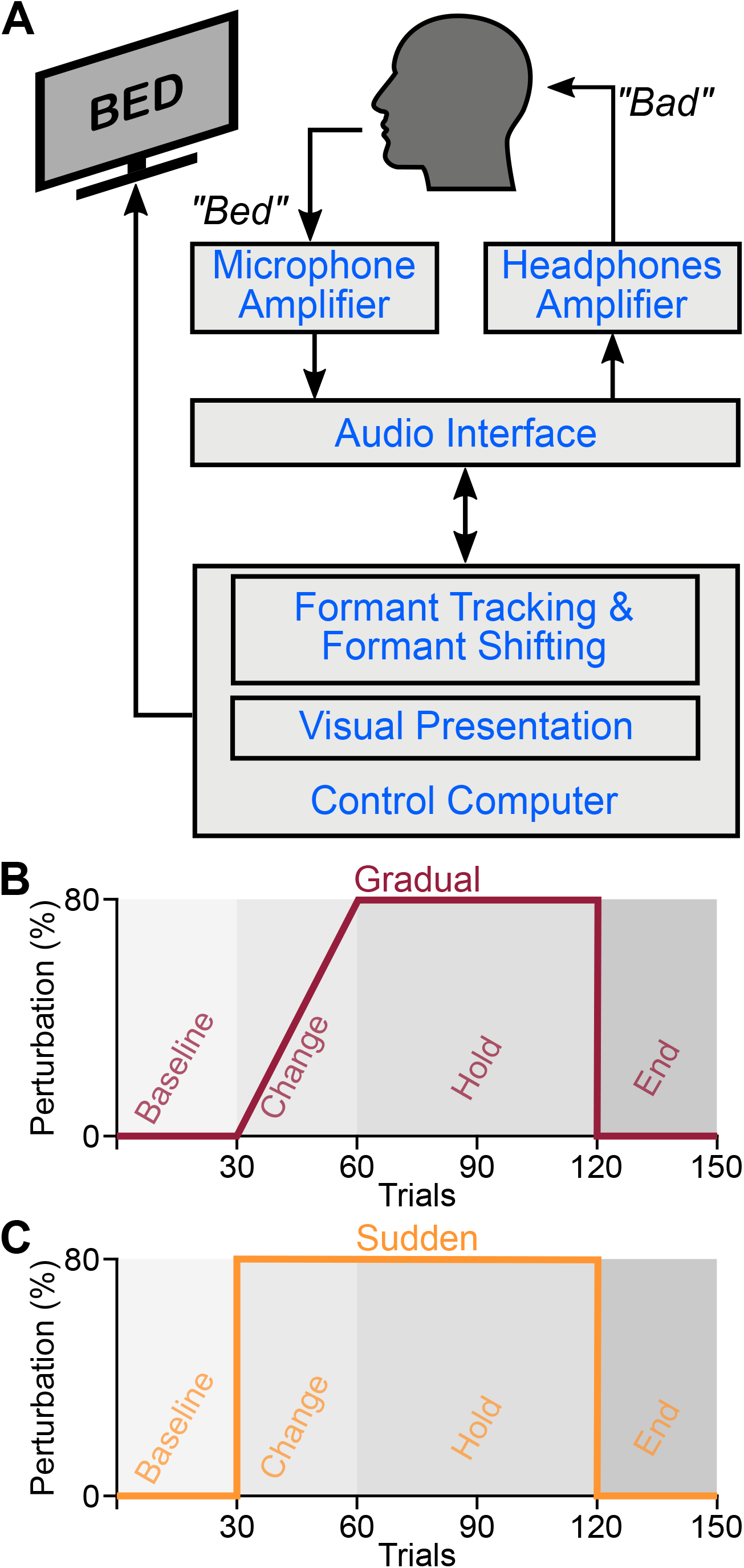
The experimental setup for applying formant perturbations (A). Each participant completed two auditory-motor adaptation blocks in which he/she received formant-shift and formant-clamp perturbations. For one group of participants, the perturbations were introduced gradually (B), and for the second group of participants, the perturbations were introduced suddenly (C). In all cases, the perturbation shifted the participant’s first formant (F1) and second formant (F2) of /ε/ toward /æ/ (increase in F1 and decrease in F2).

For auditory feedback manipulation, we used the Audapter software package (Cai, 2015). Audapter provides various routines to track and manipulate formant frequencies in near real-time. We used the following set of parameters for Audapter: sampling rate of 48 kHz, downsampling factor of 3, buffer size of 96 bytes, and a linear prediction model order of 17 for male participants and 15 for female participants (∼18 ms input-output delay). In the present study, we used two types of formant perturbations: formant-shift and formant-clamp. In the formant-shift procedure, Audapter applies a relative shift to formants such that it conserves the overall formant trajectories (e.g., 10% increase in formants). In our previous study (Daliri & Dittman, 2019), we modified Audapter’s source code to apply formant-clamp perturbations. In the formant-clamp procedure, Audapter changes formants to user-defined formant values, regardless of the original formant values (e.g., the first formant would be shifted to 600 Hz regardless of the value of the formant). Because in the formant-shift perturbations, the shifted formants (that the participant hears in the earphones) are dependent on the participant’s formants, the participant can change his/her formants to change what he/she hears in the earphones.

However, in the formant-clamp perturbations, the formants that the participant hears in the earphones are independent of the participant’s formants; thus, the participant cannot change his/her formants to change what he/she hears in the earphones. In other words, there is a close correspondence between the participant’s formants and the formants he/she hears in the earphones in the formant-shift perturbations (i.e., correspondence between motor commands and auditory feedback). However, this is not the case for the formant-clamp perturbations. Overall, the auditory feedback in the formant-shift perturbation is task-relevant, whereas the auditory feedback in the formant-clamp perturbation is task-irrelevant.

### Procedure

Participants completed three experimental tasks: one block of a practice task, one block of a pre-test task, and two blocks of an adaptation task. The entire experimental session lasted less than one hour. In the practice task, participants produced ten repetitions of consonant-vowel-consonant words (“bed,” “Ted,” and “head”). In each trial, one of the target words (in random order) appeared on the monitor for 2.5 s, followed by an intertrial interval of 1–2 s. Participants read the words aloud and received normal unperturbed auditory feedback. After each production, we provided visual feedback regarding the duration and intensity of the produced word. This task was designed to train participants to produce the target words within an intensity range of 70–80 dB SPL and a duration range of 400–600 ms.

The pre-test task was similar to the practice task, except that participants produced 30 repetitions of consonant-vowel-consonant words containing front vowels (/bɪd/, /bεd/, /bæd/). Immediately after the completion of this task, we calculated the average first formant (F1) and second formant (F2) for each production (from the middle 20% of formant trajectories estimated by Audapter). We then calculated the centroid of each vowel and visually inspected the centroids. We used the distance and angle between the centroids of /ε/ and /æ/ (in the F1-F2 coordinates) to determine the perturbation magnitude and angle for each participant. We also used F1 and F2 of the centroid of /ε/ as initial values to improve Audapter’s formant tracking algorithms.

As mentioned before, we included data from two groups of participants: a new group of participants and a group of participants from our published study (Daliri & Dittman, 2019). Each group of participants completed two blocks of the adaptation task. Each block of the adaptation task consisted of four phases in the following order: *baseline, change, hold*, and *end*. In each trial of the adaptation task, the participant produced one of the target words (“bed,” “Ted,” and “head”) while receiving unperturbed or perturbed auditory feedback. In the baseline phase (30 trials), participants received unperturbed (normal) auditory feedback. In the change phase (30 trials), participants received perturbed auditory feedback that either started in the first trial of this phase (sudden) or gradually increased over the trials of this phase until it reached the maximum perturbation level (gradual). Participants from our previous study received gradual perturbations (Figure 1B), and the new group of participants received sudden perturbations (Figure 1C). The maximum perturbation was 80% of the participant-specific ε–æ distance, such that F1 and F2 of /ε/ were shifted toward /æ/ (i.e., an increase in F1 and a decrease in F2). In the hold phase (60 trials), participants received the auditory perturbations with the maximum magnitude (80% of ε– æ). In the end phase (30 trials), participants received unperturbed (normal) auditory feedback.

Each group of participants completed two blocks of the adaptation task. We used formant-shift perturbation in one block and formant-clamp perturbation in the second block. We counterbalanced the order of the blocks within each group of participants.

### Data analysis

We used a custom-written MATLAB script to manually define the onset and offset of the vowel in each trial of the adaptation task. Trials that contained mispronunciations or formant tracking errors were excluded from the analysis. We extracted F1 and F2 trajectories starting from the onset time and ending at the offset time. Because the early portion of the vowel is primarily influenced by the feedforward control system (Daliri, 2021; Kearney et al., 2020), we averaged formant values within the first 100 ms of each vowel production to calculate the adaptive response. We designed the perturbations based on participant-specific vowel configurations such that the perturbations were parallel to the participant-specific ε–æ line (i.e., perturbation line). Similar to our previous studies (Chao et al., 2019; Daliri et al., 2020; Daliri & Dittman, 2019), we projected the average F1 and F2 of each trial to the ε–æ perturbation line (with the centroid of /ε/ as the reference point). Positive projected responses were in the direction toward /æ/. Given that the target words were in the context of /Cεd/ (“bed,” “Ted,” and “head”), the starting formant transition (from the consonant to the vowel) of the three target words were different. Therefore, we corrected the projected responses based on the average responses in the baseline phase on a word-specific basis to minimize the effects of formant transitions on the responses (Daliri, 2021; Daliri et al., 2020). Finally, to directly compare adaptive responses across participants, we divided the responses by the participant-specific ε–æ distance. We calculated these adaptive responses for the adaptation tasks and entered them into the statistical analysis.

### Statistical analysis

We performed all statistical analyses using R version 4.1.0 (R Core Team, 2021). As a dependent variable, we calculated the average adaptive responses based on (1) the 30 trials of the change phase, (2) the last 30 trials of the hold phase, (3) and the 30 trials of the end phase (see gray-shaded areas in Figure 2 A and B). The average responses in each phase and for each perturbation type and group are listed in Table 1. Because the responses were adjusted based on the average of the baseline trials, we did not enter the average responses of the baseline in the analysis. To statistically examine the adaptive responses, we used a linear mixed-effect model with perturbation group (sudden and gradual), perturbation type (formant-shift and formant-clamp), and phase (change, hold, and end phases) as fixed factors. We also entered participant as a random intercept in the linear mixed-effect model. We used the lmerTest package with Satterthwaite’s method for approximating the degrees of freedom to determine the statistical significance of the main effects and their interactions (Kuznetsova et al., 2017). For post-hoc analysis of statistically significant interactions, we used the emmeans package and Bonferroni method to correct for multiple comparisons (Lenth, 2019). Additionally, to evaluate the relationship between adaptive responses to formant-shift and formant-clamp in different phases, we used the Psych package (Revelle, 2018) to calculate Pearson’s correlation coefficients.

**Figure 2.**
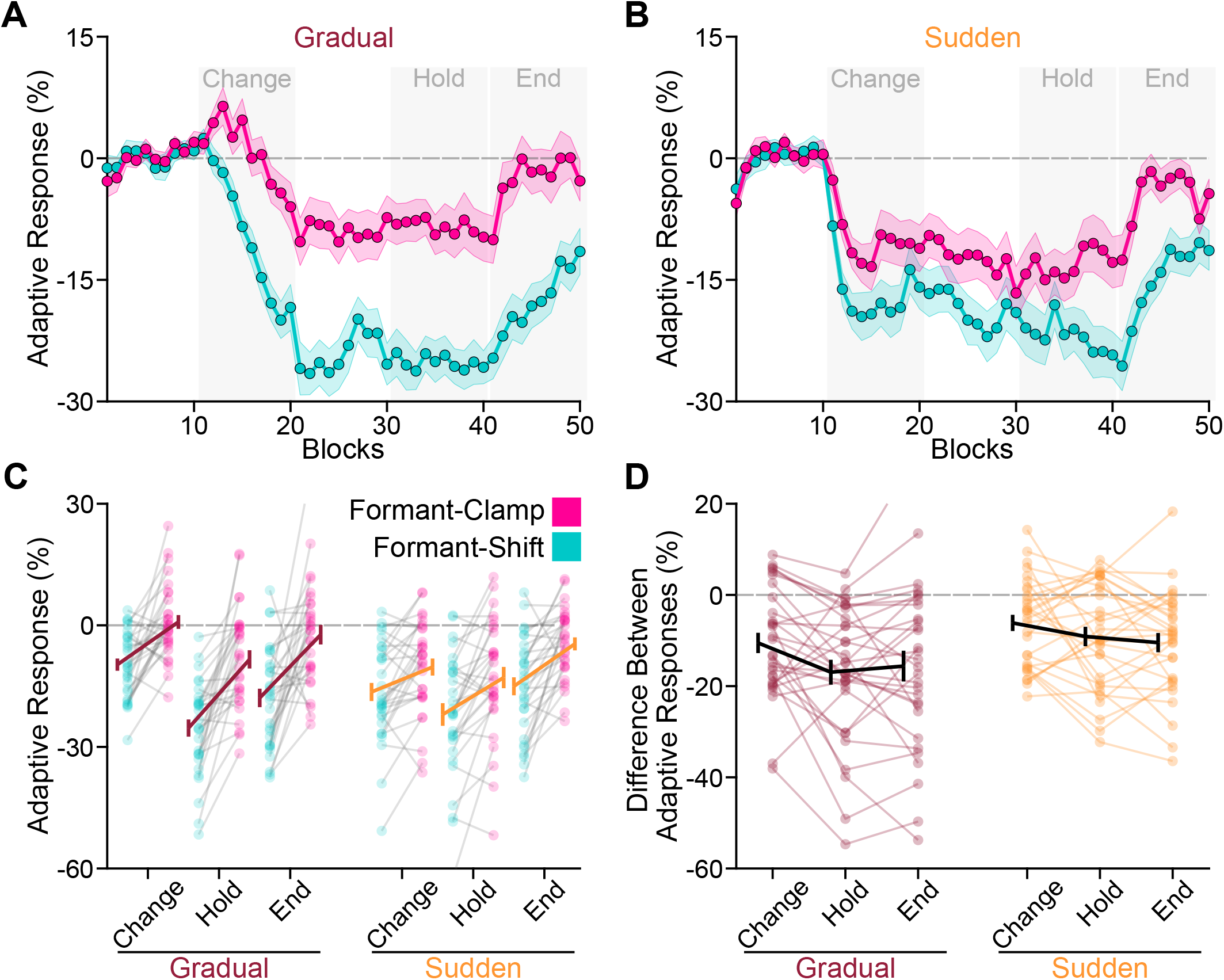
The group-average adaptive responses to formant-shift and formant clamp perturbations when perturbations were introduced gradually (A) and suddenly (B). Color-shaded areas in panels A and B correspond to the standard error of the mean. We calculated the average adaptive responses in the change, hold, and end phases (gray-shaded areas in panels A and B). Panel C shows the individual responses to different formant perturbation types (formant-shift and formant-clamp), perturbation groups (gradual and sudden), and in different phases (change, hold, and end). We found that the overall responses to formant-clamp perturbations were smaller than responses to formant-shift perturbations. However, this effect was modified by the phase factor and the group factor. The interactions indicated that the difference between adaptive responses to formant-shift and formant-clamp perturbations was the smallest when the perturbations were introduced suddenly (*p* = .031). Panel D shows the difference between responses to formant-shift and formant-clamp perturbations (responses to formant-shift perturbations minus responses to formant-clamp perturbations) for different phases and different groups. We also found that the difference between adaptive responses to formant-shift and formant-clamp perturbations was the smallest in the change phase (*p* < .016). Note that we published the data from participants of the gradual group in a previous study (Daliri & Dittman, 2019); however, in this study, we re-analyzed the data to calculate adaptive responses within 0–100 ms of the vowel onset. Error bars in panels C and D correspond to the standard error of the mean.

**Table 1.** Average adaptive responses (and standard deviations inside the parentheses) to formant-shift and formant-clamp perturbations in different phases (change, hold, and end) for both perturbation groups (gradual and sudden).

|  | Gradual |  | Sudden |  |
| --- | --- | --- | --- | --- |
|  | Formant-Shift | Formant-Clamp | Formant-Shift | Formant-Clamp |
| Change Phase | -9.70 (8.22) | 0.83 (9.29) | -16.45 (12.21) | -10.30 (11.49) |
| Hold Phase | -25.35 (11.61) | -8.40 (12.31) | -22.01 (15.29) | -12.89 (14.38) |
| End Phase | -17.92 (12.46) | -2.31 (13.21) | -15.05 (12.13) | -4.59 (8.79) |

## Results

### Difference between adaptive responses to formant-shift and formant-clamp perturbations

The primary goal of this study was to examine whether the sudden and gradual introduction of perturbations can influence adaptive responses to task-relevant (formant-shift) and task-irrelevant (formant-clamp) errors. The normalized and baseline corrected adaptative responses for the gradual and sudden groups are shown in Figure 2 A and B, respectively. We found statistically significant main effects of phase, *F*(2, 1010) = 66.951, *p* < .001, and perturbation type, *F*(1, 1010) = 327.180, *p* < .001; however, the main effect of perturbation group was not statistically significant, *F*(1, 58) = 2.036, *p* = .159. We also found phase × perturbation type interaction, *F*(2, 1010) = 6.094, *p* = .002, phase × perturbation group interaction, *F*(2, 1010) = 21.558, *p* < .001, and perturbation type × perturbation group interaction, *F*(1, 1010) = 20.846, *p* < .001. However, the phase × perturbation type × perturbation group interaction was not statistically significant, *F*(2, 1010) = 0.679, *p* = .507. The average responses in each phase and for each perturbation type and perturbation group are listed in Table 1. As shown in Figure 2 C and D, these interactions indicated that (1) the difference between adaptive responses to formant-shift and formant-clamp perturbations was the smallest when the perturbations were introduced suddenly (*p* = .031), (2) the difference between adaptive responses to formant-shift and formant-clamp perturbations was the smallest in the change phase (*p* < .016), and (3) regardless of the perturbation type, adaptive responses to gradually introduced perturbations were smaller than responses to suddenly introduced perturbations in the change phase (*p* < .001) but not in the hold and end phases (*p* > .822).

### Correlation between adaptive responses to formant-shift and formant-clamp perturbations

We conducted a series of correlational analyses to determine the relationship between adaptive responses to formant-shift and formant-clamp perturbations in different phases (change, hold, and end) and for different perturbation groups (sudden and gradual). Figure 3 shows the results of these correlational analyses. For the suddenly introduced perturbations, we found positive correlations between adaptive responses to formant-shift and formant-clamp perturbations in all three phases (change phase: *r* = .665, *p* < .001; hold phase: *r* = .691, *p* < .001; end phase: *r* = .425, *p* = .019); however, this was not the case for the gradually introduced perturbations. In other words, participants responded more similarly to formant-shift and formant-clamp perturbations when the perturbations were applied suddenly.

**Figure 3.**
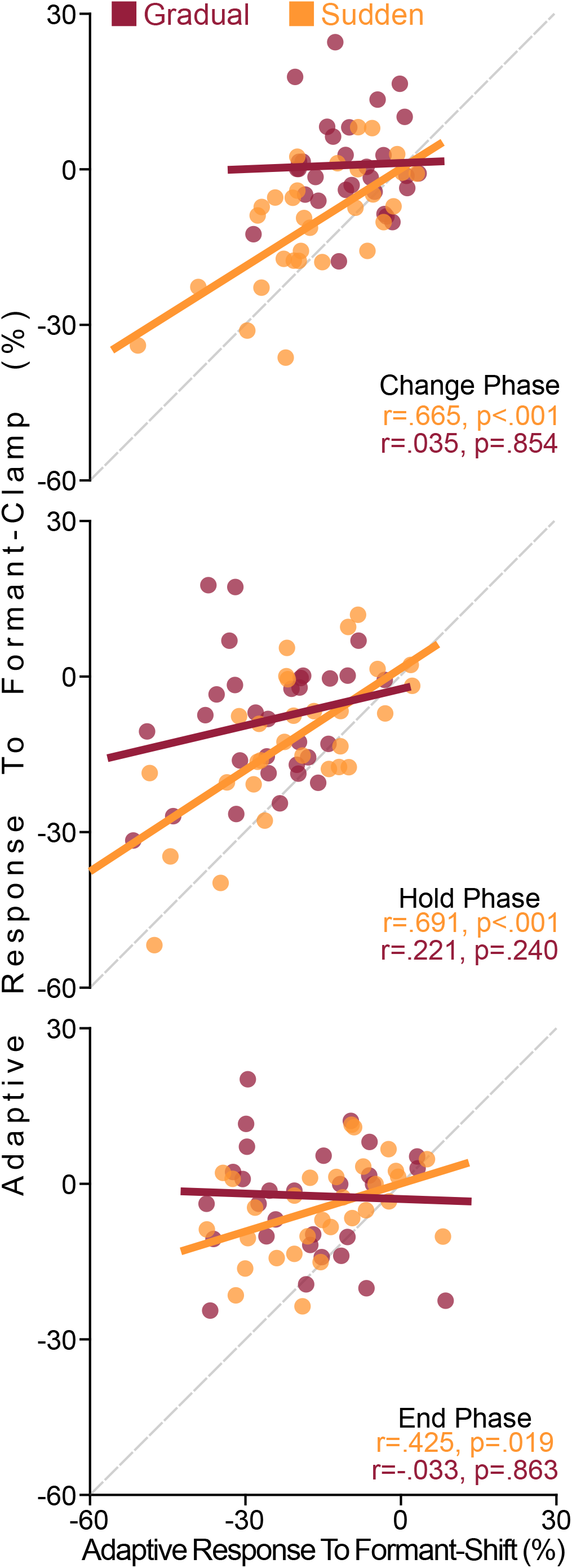
We examined the relationship between adaptive responses to formant-shift and formant-clamp perturbations in different phases (change, hold, and end) and for different perturbation groups (sudden and gradual). We found positive correlations between responses to formant-shift and formant-clamp perturbations in all three phases of the suddenly introduced perturbations; however, this was not the case for the gradually introduced perturbations. The gray dashed line is the identity line (i.e., a line with the slope of 1).

## Discussion

The primary goal of the present study was to determine how the SMS evaluates and responds to task-relevant vs. task-irrelevant errors that are introduced suddenly and gradually. In a previous study (Daliri & Dittman, 2019), we developed an experimental procedure to directly manipulate the relevance of errors (i.e., formant-clamp). In the current study, we used an auditory-motor adaptation paradigm in which we applied formant-shift perturbations (task-relevant errors) or formant-clamp perturbations (task-irrelevant errors). We compared adaptive responses to suddenly introduced formant-shift and formant-clamp perturbations (data from the current study) with responses to gradually introduced formant-shift and formant-clamp perturbations (data from the previous study). Suddenly introduced perturbations are more noticeable than gradually introduced perturbations; thus, at least initially, the SMS may evaluate suddenly introduced errors as less task-relevant errors. We hypothesized that the SMS responds to task-relevant errors (induced by formant-shift perturbations) and task-irrelevant errors (induced by formant-clamp perturbations) more differently when the errors are introduced gradually.

We found that (1) the difference between adaptive responses to formant-shift and formant-clamp perturbations was the smallest when the perturbations were introduced suddenly, and (2) adaptive responses to formant-shift perturbations positively correlated with responses to formant-clamp perturbations for suddenly introduced perturbations but not gradually introduced perturbations. These results indicated that the SMS evaluates errors and uses its evaluations to determine its adaptive responses. Generally, suddenly introduced errors are more pronounced, and the brain is more likely to assign suddenly introduced errors to external sources (Berniker & Körding, 2008; Krakauer et al., 2019; Shadmehr et al., 2010). Therefore, the SMS may evaluate suddenly introduced formant-shift perturbations as less task-relevant and integrate them like task-irrelevant errors induced by formant-clamp perturbations. While this interpretation can explain within-subject analysis (i.e., responses to formant-shift vs. formant-clamp perturbations in each group), it does not explain the results of between-subject analyses. Specifically, if the SMS evaluates suddenly introduced errors as less task-relevant, the responses to sudden perturbations should be smaller than responses to gradual perturbations. However, our between-subject analysis showed that responses to gradual perturbations were like sudden perturbations (except in the change phase) for formant-shift and formant-clamp perturbations. The results for gradually vs. suddenly introduced formant-shift perturbations are consistent with the results of previous studies (Kearney et al., 2020; Kim et al., 2020; MacDonald et al., 2010). These results may be related to the fact that the within-subject analysis is more sensitive to detect potential effects (for a review, see Charness et al., 2012) of the sudden and gradual introduction of perturbations. Future studies can examine this issue by designing experiments in which each participant experiences gradually and suddenly introduced formant-shift and formant-clamp perturbations. Overall, our results indicated that how errors are introduced modulates the SMS’s evaluation of the relevance of the errors and the SMS’s adaptive responses to the errors.

We also found that (1) adaptive responses to gradual perturbations were smaller than responses to sudden perturbations in the change phase, and (2) the difference between adaptive responses to formant-shift and formant-clamp perturbations was the smallest in the change phase. We attribute both results to the introduction of the perturbations. The adaptive responses to the gradual perturbations gradually increased throughout the change phase (30 trials) until they reached a stable level at the end of the change phase. However, adaptive responses to the sudden perturbations increased rapidly, and after ∼9 trials, they reached a relatively stable level. It should be noted that these patterns of responses were similar for both formant-shift and formant-clamp perturbations. Because we averaged responses in all 30 trials of the change phase, the overall responses were larger for the sudden perturbations than the gradual perturbations. Additionally, because the overall responses to formant-shift and formant-clamp perturbations are smaller in the change phase (compared with responses in other phases), the difference between these responses is also smaller. These results may indicate that, although the SMS evaluates formant-shift and formant-clamp perturbations differently, it uses similar procedures to respond to the errors. One limitation of the current study is that the maximum perturbation was 80% of the participant-specific ε–æ distance; therefore, it is unclear whether the perturbation magnitude differently modulates adaptive responses to formant-shift and formant-clamp perturbations. Future studies can determine the effects of perturbation magnitude by examining adaptive responses to formant-shift and formant-clamp perturbations with various magnitudes.

Our results have important theoretical and clinical implications. Most models of speech posit that the SMS calculates prediction errors and uses the errors to develop adaptive responses (Daliri, 2021; Hickok, 2012; Houde & Nagarajan, 2011; Kearney et al., 2020; Parrell et al., 2019). However, as our results indicated, the SMS evaluates the relevance of errors and uses its evaluation to modulate its adaptive responses to the experienced errors. Recently, we developed a state-space model to estimate the SMS’s sensitivity to errors and predict the SMS’s responses to various errors (Daliri, 2021). Like other models, our model used the calculated error in each trial of an adaptation task to generate a response (i.e., multiplying the calculated error by error sensitivity). Given the results of this study, we suggest that a future version of the model can include an error evaluation stage in which the SMS assigns a weight to the error based on the relevance of the error (e.g., a weight between 0 and 1, with 1 being the highest level of task-relevance). Then, the adaptive responses can be calculated based on the experienced errors, the sensitivity to the errors, and the relevance of the errors. Our results may also have clinical implications. For example, we have previously shown that adaptive responses to auditory perturbations are significantly small (or nonexistent) for adults who stutter (Daliri et al., 2017; Daliri & Max, 2018; Kim et al., 2020; Max & Daliri, 2019). Previously, we interpreted the small adaptive responses of people who stutter as lower sensitivity to errors. However, based on the current results, one could speculate that adults who stutter may evaluate errors as less task-relevant (i.e., assign a lower weight to the experienced errors). Overall, our results can be used to modify current theories of speech production and theories of disorders of speech production.

In summary, we examined the SMS’s adaptive responses to task-relevant vs. task-irrelevant errors when the errors are introduced gradually and suddenly. We used an experimental procedure to directly manipulate the relevance of errors (i.e., formant-clamp). Two groups of participants completed auditory-motor adaptation tasks; the first group received gradually introduced formant-shift and formant-clamp perturbations; the second group received suddenly introduced formant-shift and formant-clamp perturbations. We found that (1) the difference between adaptive responses to formant-shift and formant-clamp perturbations was the smallest when the perturbations were introduced suddenly, and (2) adaptive responses to formant-shift perturbations positively correlated with responses to formant-clamp perturbations for the sudden but not the gradual perturbations. These results indicated that the SMS evaluates the relevance of errors and uses its evaluation to modulate the magnitude of its adaptive responses to errors.

## Acknowledgments

This work was supported by a grant from the National Institutes of Health awarded to A. Daliri (R21 DC017563). We thank Kimiya Kasraeian and Sreelakshmi Gurrala for their contributions to participant recruitment for this project.

